# *Trem2* R47H mutation shows mild, but functionally divergent alterations in microglial phenotypes compared to *Trem2* deficiency in aged *App^NL-F^* knock-in mice

**DOI:** 10.64898/2026.05.18.724708

**Authors:** Keiro Shirotani, Daisuke Hatta, Kaori Watanabe, Takashi Saito, Takaomi C. Saido, Nobuhisa Iwata

**Affiliations:** Department of Genome-based Drug Discovery, Graduate School of Biomedical Sciences, Nagasaki University, Nagasaki 852-8521, Japan; Leading Medical Research Core Unit, Graduate School of Biomedical Sciences, Nagasaki University, Nagasaki 852-8521, Japan; Department of Neuropathology, Graduate School of Medicine, The University of Tokyo, Tokyo 113-0033, Japan; Laboratory for Proteolytic Neuroscience, RIKEN Center for Brain Science, Saitama 351-0198, Japan

**Keywords:** Alzheimer’s disease, risk gene, TREM2, microglia, amyloid β, dystrophic neurite, gene expression

## Abstract

The TREM2 R47H variant increases the risk of Alzheimer’s disease (AD), yet its functional impact in aged mouse models remains incompletely understood. We generated a humanized *Trem2* R47H knock-in (KI) line on the *App^NL-F^* background and compared it with a *Trem2* knockout (KO) line to assess the degree of TREM2 functional impairment. Accumulation of amyloid β 42 and formation of dystrophic neurites were increased in *Trem2* KO mice but not in *Trem2* R47H KI mice at 18 or 24 months. qPCR and transcriptomic analyses revealed *Trem2* KO mice showed deficits in upregulation of microglial genes while *Trem2* R47H KI mice showed a response similar to control mice. Differential gene expression analysis identified altered expressions of genes responsible for ER stress/unfolded protein response and intracellular signalling in *Trem2* R47H KI mice. Among the differentially expressed genes, *Pmel* and *Gpnmb* were or tended to be downregulated in *Trem2* R47H KI as well as in *Trem2* KO mice indicating their involvement in AD pathogenesis. These results clearly indicate that the TREM2 R47H variant confers a mild, rather than null, effect on microglial alterations during AD development and that *Trem2* R47H KI mice should be used to understand pathological mechanism elicited by TREM2. Further identification and characterization of genes differentially expressed in *Trem2* R47H KI mice will provide important insights into how the TREM2 risk variant modulates Alzheimer’s disease–related pathology.

**Highlights:**

- Exon2-humanized *Trem2* R47H knock-in mice are established, which will serve as a platform to study the role of TREM2 in Alzheimer’s disease development.
- *Trem2* knockout mice exhibit deficits in clearance of highly aggregated Aβ42, suppression of dystrophic neurites and regulation of microglial genes in *App^NL-F^* mice, whereas *Trem2* R47H knock-in mice do not.
- RNA-seq reveals transcriptional profiles of *Trem2* R47H knock-in mice
- qPCR confirms that *Gpnmb* and *Pmel* are or tended to be downregulated in *Trem2* R47H knock-in mice.
- Findings demonstrate that TREM2 R47H is hypomorphic rather than loss of function.

## 1. Introduction

Alzheimer’s disease (AD) is the most prevalent dementia worldwide. Although anti-amyloid-β (Aβ) antibodies, lecanemab and donanemab, have been approved as disease-modifying drugs in the USA, Japan, and other countries, drugs targeting other molecules should be developed for more effective cure in an economical manner. Genetic studies have identified several AD-associated risk genes expressed in microglia (Bertram et al., 2008; Hollingworth et al., 2011; Lambert et al., 2009; Naj et al., 2011), among which triggering receptor expressed on myeloid cells 2 (TREM2) has the highest odds ratio for AD (R. Guerreiro et al., 2013; Jonsson et al., 2013) . TREM2 is a type I transmembrane glycoprotein expressed on the surface of microglia and transduces intracellular signals, such as cytokine production, migration, proliferation, cell survival, phagocytosis, compaction of amyloid plaques and synapse elimination (Filipello et al., 2018; Jay et al., 2017, 2015; Mazaheri et al., 2017; Takahashi et al., 2005; Ulrich et al., 2014; Wang et al., 2016, 2015; Yuan et al., 2016). Moreover, homozygous mutations in *TREM2* have been found in Nasu-Hakola disease and frontotemporal dementia-like syndromes (Chouery et al., 2008; R. J. Guerreiro et al., 2013; Paloneva et al., 2002) indicating that these diseases are developed by a loss of function of TREM2.

To understand the mechanism by which TREM2 accelerates AD development, *Trem2* gene was first deleted in amyloid-β precursor protein (*APP*) transgenic (tg) mice (5xFAD, APPPS1, or PS2APP mice). *Trem2* knockout (KO) impairs microglial clustering (Jay et al., 2015; Meilandt et al., 2020; Wang et al., 2015) and barrier function around amyloid plaques (Wang et al., 2016; Yuan et al., 2016), enhances Aβ accumulation (Jay et al., 2017; Wang et al., 2015) and seeding of Aβ plaques (Parhizkar et al., 2019), and exacerbates the pathology of dystrophic neurites (Meilandt et al., 2020; Wang et al., 2016; Yuan et al., 2016). However, the effects of *Trem2* deletion on Aβ accumulation are contradictory depending on the mouse model, age and sex. In *Trem2^KO^*x 5xFAD mice, Aβ amounts in the cerebral cortex and hippocampus are unchanged at 4 months and increased at 8 months (Wang et al., 2015). In *Trem2^KO^*x APPPS1 mice, amyloid plaque load is temporally reduced at 2–4 months but increased at 8 months (Jay et al., 2017, 2015). In female *Trem2^KO^*x PS2APP mice Aβ burden is not altered at 4 months and increased at 6–7 months but decreased at 12 months, while it is not changed at 4–7 months but decreased at 19–22 months in male (Meilandt et al., 2020).

Thereafter, *Trem2* R47H knock-in (KI) mice were produced but most of the mice expressed low amounts of full-length *Trem2* R47H mRNA due to cryptic mRNA splicing in mouse exon2 containing R47H, which is not observed in human exon2 (Cheng-Hathaway et al., 2018; Xiang et al., 2018). Human *TREM2* R47H cDNA was then knocked-in on the mouse *Trem2* locus (Sayed et al., 2021), but the *TREM2^R47H/+^* mice did not enhance the accumulation of Aβ in *App^NL-G-F^* mice (Das et al., 2023), *App* KI mouse model with aggressive Aβ accumulation as *APP* tg mice (Saito et al., 2014). Tran et al. successfully established *Trem2* R47H KI mice by carefully avoiding cryptic splicing and crossed them with 5xFAD mice (Tran et al., 2023). The *Trem2^R47H^* x 5xFAD mice, however, showed an increase, decrease, or no change in Aβ accumulation depending on sex, age and brain region. The number and size of microglia around amyloid plaques were reduced and dystrophic neurite formation were enhanced at 4 months in the *Trem2^R47H^*x 5xFAD mice but these phenotypes diminished at 12 months. Taken together, *Trem2* R47H KI mice have shown only a mild phenotype in model mice with aggressive Aβ accumulation.

The *App^NL-F^* KI model provides a physiologically relevant platform for evaluating microglial responses and Aβ accumulation without overexpression artifacts of APP (Saito et al., 2014). Aβ plaques in this model begin at 6 months and continue to increase until at least 24 months (Saito et al., 2014). This slower Aβ progression compared to that in *APP* tg and *App^NL-G-F^* mice allows for the investigation of microglial function in aged animals before Aβ accumulation reaches a plateau. To address the impact of TREM2 R47H under physiologically relevant conditions and to elucidate the impact of varying levels of TREM2 impairment on AD pathology, we generated *Trem2* R47H KI and *Trem2* KO lines on the *App^NL-F^*background and compared these phenotypes in aged mice. This comparative approach enables the evaluation of whether the R47H variant behaves as a hypomorphic allele and clarifies the degree of TREM2 function required to maintain microglial activation and suppress neuritic damage during late-stage Aβ accumulation. We found that *Trem2* KO mice displayed an increase of highly aggregated Aβ42, enhanced dystrophic neurite formation, and decrease of microglial gene expression, whereas *Trem2* R47H KI mice did not. Transcriptome analysis identified genes related to ER stress/unfolded protein response and intracellular signalling as differentially expressed genes in *Trem2* R47H KI mice compared with control. Further characterization of these genes will contribute to the identification of novel mechanisms underlying AD pathogenesis.

## 2. Materials and methods

### 2.1. Animals

All animal experiments were approved by Nagasaki University Animal Care and Use Committee. *App* knock-in mice *App^NL–F^* were as described (Saito et al., 2014) and *Trem2* KO and KI mice were generated in this study. They received food and water ad libitum and were maintained on a 12/12-h light–dark cycle (lights on at 08:00, off at 20:00). Homozygous female mice were analyzed at the indicated ages. Mice were deeply anesthetized by intraperitoneal injection of 6 mL/kg mixture of 0.5 mg/mL butorphanol tartrate (Meiji Seika Pharma, Tokyo, Japan), 0.15 mg/mL medetomidine hydrochloride (Kyoritsu Seiyaku, Tokyo, Japan), and 0.8 mg/mL midazolam (Sandoz K.K., Tokyo, Japan), and transcardially perfused with PBS. One brain hemisphere was immersion-fixed in 4% paraformaldehyde in PBS and cryoprotected in 30% sucrose for immunohistochemical analyses. The cerebrocortical and hippocampal tissues from the other hemisphere were subdissected and frozen at –80°C for biochemical assays.

### 2.2. Generation of Trem2 KO mice

*Trem2* KO mice were generated by CRISPR/Cas9-assisted gene targeting at Research Center for Advanced Genomics, Nagasaki University. Briefly, the pronuclei of fertilized oocytes obtained from superovulated C57BL/6N females were microinjected with Cas9 protein (IDT, San Diego, CA, USA), *Trem2* crRNA in exon2 corresponding to gaacctccaagccggtgacg<u>cgg</u> where underlined nucleotides indicate PAM site, and tracrRNA (IDT). crRNAs with minimal off-target effects were selected using the IDT and UCSC browsers. Genomic DNAs from offspring were isolated from the tissues and the loci were amplified by PCR using the primer sets (attaaccatcttgcacacgc and aacccttgccaggcacctac) and sequenced. To exclude off-target effects, the knockout F0 mice were backcrossed with the C57BL/6J genetic background.

### 2.3. Generation of exon2-humanized Trem2^hE2WT^ and Trem2^hE2R47H^ knock-in mice

The pronuclei of fertilized oocytes obtained from superovulated C57BL/6N females were microinjected with Cas9 protein (IDT), the Trem2 crRNA described above, tracrRNA (IDT), and 586 nt single stranded DNA (ssDNA) (IDT) for recombination. The ssDNA sequence is ccagaacttatcctaatgaccatgcacacgctatgctccctgcactcctggaactgcttccaagcaagtggctgtctcctctgcagAGCTGTC CGGAGCC<u>GACTACAAAGACCATGACGGTGATTATAAAGATCATGACATCGATTACAAG</u> <u>GATGACGATGACAAG</u>CACAACACCACAGTGTTCCAGGGCGTGGCGGGCCAGTCCCTGC AGGTGTCTTGCCCCTATGACTCCATGAAGCACTGGGGGAGGC**<u>A</u>**CAAGGCCTGGTGCCGC CA<u>A</u>CTGGGAGAGAAGGGCCCATGCCAGCGTGTGGTCAGCACGCACAACTTGTGGCTGC TGTCCTTCCTGAGGAGGTGGAATGGGAGCACAGCCATCACAGACGATACCCTGGGTGG CACTCTCACCATTACGCTGCGGAATCTACAACCCCATGATGC<u>C</u>GGTCTCTACCAGTGCC AGAGCCTCCATGGCAGTGAGGCTGACACCCTCAGGAAGGTCCTGGTGGAGGTGCTGGCAGgtgagtacacagtagctgggtgcacctttggctggtccttgtgccaggtcttattgttgtgggactttttcagggggacataa representing small letters as the mouse intron sequence, capital letters as the human exon2 sequence, and underlined capital letters as the 66 nt 3 x FLAG sequence after the signal peptide. The substitution from G to A for the R47H mutation is marked by an underlined bold capital letter. Additional silent mutations at a possible unwanted splice site (G to <u>A</u>) (Xiang et al., 2018) and at a corresponding PAM site in human exon2 which may be reablated by Cas9 (G to <u>C</u>), are underlined. The Trem2 loci of offspring were amplified by PCR, and the *Trem2^hE2R47H (w/3xFLAG)^* allele whose exon2 was replaced by human exon2 with 3xFLAG after the signal peptide was identified by sequencing.

Next, the pronuclei of fertilized oocytes from *Trem2^hE2R47H (w/3xFLAG)^* knock-in mice were used for microinjection of Cas9 protein, Trem2 crRNA in exon2 corresponding to cctatgactccatgaagcac<u>tgg</u> where underlined are PAM site, tracrRNA (IDT), and 84 nt ssDNA (IDT) to create *Trem2^hE2WT (w/3xFLAG)^* whose R47H in human exon2 was reverted to wild-type sequence (A to G). The ssDNA sequence is CCAGTCCCTGCAGGTGTCTTGCCCCTATGACTC<u>A</u>ATGAA<u>A</u>CA<u>T</u>TGGGGGAGGC**<u>G</u>**CAAGGCCTGGTGCCGCCAACTGGGAGAGAA, where the A to G substitution for wild-type is marked by underlined bold letter, and three additional silent mutations to avoid reablation at the PAM site are underlined. The *Trem2* loci of offspring were amplified by PCR, and the *Trem2^hE2WT (w/3xFLAG)^* allele was identified by sequencing.

Finally, to delete the 3xFLAG sequence from *Trem2^hE2WT (w/3xFLAG)^* and *Trem2^hE2R47H (w/3xFLAG)^* alleles, the pronuclei of fertilized oocytes of the mice were microinjected with Cas9 protein (IDT), two *Trem2* crRNAs in exon2 corresponding to gatctttataatcaccgtca<u>tgg</u> and agatcatgacatcgattaca<u>agg</u> where underlined are PAM sequence, tracrRNA (IDT) and 145 nt ssDNA (IDT). The ssDNA sequence is CATGCACACGCTATGCTCCCTGCACTCCTGGAACTGCTTCCAAGCAAGTGGCTGTCTCCT CTGCAGAGCTGTCCGGAGCCCACAACACCACAGTGTTCCAGGGCGTGGCGGGCCAGTCCCTGCAGGTGTCTTGCCCCTATGACTC, which lacks the 3xFLAG sequence. The *Trem2* loci of offspring were amplified by PCR, and *Trem2^hE2WT^* and *Trem2^hE2R47H^*alleles without 3xFLAG were identified by sequencing. To avoid off-target effects, the KI mice were then backcrossed to a C57BL6/J background.

### 2.4. Mouse genotyping procedure

Genotyping for *App^NL-F^* was performed as previously described (Saito et al., 2014). The primers used for *Trem2* genotyping were as follows: forward primer gcaagtggctgtctcctctg and reverse primer cacccagctactgtgtactc. Amplified PCR products were electrophoresed in 1.5% agarose gel to distinguish between *Trem2^KO^* and wt (mouse wild-type) by their mobility. To distinguish between *Trem2^hE2WT^*, *Trem2^hE2R47H^* and wt, the PCR products were digested with the restriction enzymes *Bgl*I (TOYOBO, Shiga, Japan) or *Hha*I (Takara Bio, Osaka, Japan) and electrophoresed. Genotypes were determined as *Bgl*I cuts *Trem2^hE2WT^*and *Trem2^hE2R47H^* while *Hha*I cuts *Trem2^hE2WT^*.

### 2.5. Preparation of mouse primary microglia

The cerebral cortices of newborn mice (P0–P1) were dissected, and the meninges were carefully removed. The cortices were trypsinized for 10 min at 37°C with gentle agitation and incubated for 30 s at room temperature with 0.05% DNase I (Roche). Trypsinization was stopped using fetal bovine serum (FBS), and cells were collected by centrifugation, triturated by pipetting, and seeded in poly-L-lysine coated culture flasks in DMEM (Fujifilm Wako, Osaka, Japan) supplemented with 10% FBS and penicillin/streptomycin and cultured in a humidified atmosphere of 5% CO_2_ at 37°C. The media were replaced with DMEM on day 4 and then with DMEM containing 20 ng/ml mouse macrophage-colony stimulating factor (Fujifilm Wako) on day 7. The mixed glial cells were cultured for additional 7 days. After shaking the culture flask for 5 min at 100 rpm, the culture media containing microglial cells were centrifuged and subjected to immunoblot analysis.

### 2.6. Immunoblot analysis

Immunoblot analysis was performed as previously described (Shirotani et al., 2023, 2022, 2019). Briefly, cell lysates were extracted with lysis buffer (1% Triton-X 100, protease inhibitor cocktail (Complete^TM^, Roche Diagnostics, Indianapolis, IN, USA), and 700 ng/ml pepstatin A (Peptide Institute, Osaka, Japan) in phosphate-buffered saline). Aliquots were electrophoresed on 10% polyacrylamide sodium dodecyl sulfate-gel and blotted onto a nitrocellulose membrane (Protran BA 83, Cytiva, Marlborough, MA, USA). The membrane was probed with anti-mouse Trem2 antibodies (Cell Signaling Technology, Danvers, MA, USA), followed by HRP-conjugated secondary antibodies IgG (Cell Signaling Technology). Immunoreactive bands were visualized using ImmunoStar LD (Fujifilm Wako) and detected on a LAS-4000 mini imager (Fujifilm Wako).

### 2.7. Aβ42 ELISA

Aβ extraction from brain tissues was performed as described previously (Iwata et al., 2024). Mouse cerebral cortices were homogenized in buffer A (50 mM Tris–HCl, pH 7.6, 150 mM NaCl, and protease inhibitor cocktail) using an RW20 digital (IKA Japan, Osaka, Japan). The homogenized samples were centrifuged at 200,000 *x g* for 20 min at 4°C. The supernatant was excluded from the analysis because it contained tiny amounts of Aβ42 in *App^NL-F^*mice. The pellet was dissolved in 6 M guanidine hydrochloride (GuHCl) buffer and sonicated at 25°C for 1 min. After incubation at room temperature for 1 h, the sample was centrifuged at 200,000 *x g* for 20 min at 25°C, and the supernatant was collected as a GuHCl fraction. The pellet was subsequently dissolved in 100% formic acid, agitated at 25°C for 30 min, and centrifuged at 200,000 *x g* for 20 min at 25°C. The supernatant was dried using speed vac and dissolved in DMSO, followed by sonication for 30 s and agitation for 10 min. Sandwich ELISA by BAN50 as the capture antibody and BC05 as the detection antibody to quantify Aβ42 was described previously (Iwata et al., 2001).

### 2.8. Immunohistochemistry

The fixed brain hemispheres were embedded in OCT compound FSC 22 Blue (Leica, Wetzlar, Germany) and sectioned at 25 μm using a Leica CM1950 freezing microtome (Leica). The sections were adhered to micro slide glass FRC-01 (Matsunami Glass, Osaka, Japan) and incubated in PBS containing 0.25% Triton X-100 for 20 min at room temperature. Endogenous peroxidases were quenched with 0.3% hydrogen peroxide in methanol for 30 min, and the sections were blocked with Tris-NaCl-blocking buffer FP1012 (PerkinElmer, Waltham, MA, USA). The sections were incubated with the following primary antibodies overnight at 4°C: anti-Iba1 antibody (Fujifilm Wako), anti-Aβ antibody 82E1 (IBL, Osaka, Japan) or N1D (Saido et al., 1995), and anti-Lamp-1 antibody 1D4B (Santa Cruz Biotechnology, Dallas, TX, USA). Signals were visualized using the appropriate secondary antibodies and tyramide signal amplification system (Revvity, Waltham, MA, USA; Proteintech Rosemont, IL, USA; AAT Bioquest, Pleasanton, CA, USA; Dako, Glostrup, Denmark; Cell Signaling Technology). The sections were then mounted in mounting medium TA-030-FM (Thermo Fisher Scientific, Waltham, MA, USA) and covered with micro cover glasses (Matsunami Glass). *z*-Stack images were collected at 20× or 100× objectives under a fluorescence microscope BZ-9000 (KEYENCE, Osaka, Japan), and maximum intensity projection images were created. For the quantification of dystrophic neurites, images were acquired at 20× objective on a digital slide scanner Nanozoomer S60 (Hamamatsu Photonics, Shizuoka, Japan). All image acquisition settings were maintained in each experiment.

### 2.9. Image analysis

Quantification analysis of dystrophic neurites was performed in the cerebral cortex and hippocampus using Halo software (Indica Labs, Albuquerque, NM, YSA) according to previous studies (Cheng-Hathaway et al., 2018; Cuddy et al., 2022; Sadleir et al., 2022a; Yin et al., 2023). Briefly, the threshold of the fluorescence signal was determined to isolate the signals of interest by optimizing the signal-to-noise ratio and eliminating nonspecific background signals. Amyloid plaques in the cortex and hippocampus ranging from 70 to 400, 400 to 800, and more than 800 μm^2^ (5 to 20 plaques in each category area from each mouse) were chosen. The areas of amyloid plaques and the surrounding anti-Lamp-1 positive dystrophic neurites were measured by the software and the ratio of the dystrophic neurite to amyloid plaque area from each plaque were calculated and averaged in each category area for each animal.

### 2.10. RNA isolation and quantitative reverse transcription PCR (qPCR)

Total RNA was extracted from the cortices of 18-month-old female mice using Nucleospin RNA (Takara Bio). First-strand cDNA synthesis and qPCR were conducted using One Step Primescript RT-PCR Kit (Takara Bio) in a Lightcycler96 system (Roche, Basel, Switzerland) with the following profile: 5 min at 42°C and 50 cycles of 5 s at 95°C and 20 s at 60°C. Relative quantification of RNA expression was carried out by normalization to the expression of glyceraldehyde 3-phosphate dehydrogenase (*Gapdh*). The nucleotide sequences of the forward/reverse primers/probes were as follows: *Aif1*; agctgaagagattaattagagaggtg/tctcctcatacatcagaatcattctc/tggctccgaggagacgttcagctactc, *Casr*; tctccagagaggtgcccttc/cacaggcactcgcatctgtc/actcgccgtcaggacactccacac, *Cd68*; tggacagcttacctttggattca/ctgtgctgcctgtgggaag/acaggacctacatcagagcccgagtacagt, *Gapdh*; caagctcatttcctggtatgacaa/ttgggatagggcctctcttgc/tggtggacctcatggcctacatggcc, *Gpnmb*; tgtgcctgctgtctgtgag/ttgggatagggcctctcttgc/gggagtctgggtctttgcc, *Pmel*; ggctcccttgcttgtaggta/ttgggatagggcctctcttgc/cctgcttcttaagtctatgcctatg, *Tmem119*; ccttcacccagagctggttcc/ttgggatagggcctctcttgc/cggctacatcctccaggaagg, *Trem2*; tgcgttctcctgagcaagtttc/cgtacctccgggtccagtg/ccacagtccagccctctgaccacagg, *Tyrobp*; aagggacccggaaacaacac/ctctgtgtgttgaggtcactgtata/aagctcctgataaggcgactcagtctcagc.

### 2.11. RNA sequencing, differential gene expression and GO analysis

RNA sequencing (RNA-seq) analysis was outsourced to Cell Innovator (Fukuoka, Japan). Briefly, total RNAs were quantified by an ND-1000 spectrophotometer (NanoDrop Technologies, Wilmington, DE), and the quality was confirmed with a Tapestation (Agilent, Santa Clara, CA, USA). The sequencing libraries were prepared from 200 ng of total RNA with MGIEasy rRNA Depletion Kit and MGIEasy Fast RNA Library Prep Set (MGI Tech, Shenzhen, China) according to the manufacturer’s instructions. The libraries were sequenced on the DNBSEQ-G400 FAST Sequencer (MGI Tech) with paired-end 150 nt strategy. All sequencing reads were trimmed of low-quality bases and adapters with Trimmomatic (v.0.38) (Bolger 2014). Trimmed reads were mapped to the transcript in the mouse GRCm39 using the Bowtie2 aligner within RSEM (Langmead and Salzberg, 2012; Li and Dewey, 2011). The Abundance estimation of genes and isoforms with RSEM generated basic count data (expected counts). Raw RNA-Seq data were deposited in NCBI GEO under accession number GSE328142. We used edgeR program (Robinson et al., 2009) to detect differentially expressed genes (DEGs). Normalized counts per million values, log fold-changes (log_2_FC), and p-values were obtained from the gene-level raw counts. We established the criteria for DEGs as p-value <0.05.

To determine significantly over-represented Gene Ontology (GO) categories and enriched pathways, functional enrichment analysis was performed using the Database for Annotation, Visualization, and Integrated Discovery (DAVID) (http://david.abcc.ncifcrf.gov/) (Huang et al., 2009). Statistical significance was evaluated using a modified Fisher’s exact test (the EASE score). Heatmaps were generated using the MultiExperiment Viewer (MeV). Normalized gene expression values were visualized after row-wise Z-score normalization.

### 2.12. Statistical analyses

Quantitative data of animal experiments are expressed as mean ± standard error. Statistical analyses were done using SigmaPlot software ver.14.0 (Systat Software Inc., San Jose, CA, USA). For comparisons of the means between the two groups, statistical analysis was performed using Student’s t-test. The effects of two factors (genotype and amyloid plaque area) on the dystrophic neurite/amyloid ratio and their interactions were analyzed by two-way ANOVA, followed by a *post hoc* Student–Newman–Keuls multiple comparison test. Differences were considered significant when *p*-values were less than 0.05.

## 3. Results

### 3.1. Generation of Trem2 knockout and Trem2 R47H knock-in mice

We generated *Trem2* KO (*Trem2^KO^*) and *Trem2* R47H KI mice by CRISPR/Cas9 gene editing with crRNA targeted to mouse exon2. *Trem2^KO^* mice lack 35 nt in exon2 (Fig. 1A) and its premature protein product has the first 65 amino acids with additional frameshifted 4 amino acids. As expected, Trem2 protein was not detected in primary microglia from *Trem2^KO^* mice (Fig. 1B lane 2). To generate *Trem2* R47H KI mice, DNA oligonucleotides corresponding to the entire human *TREM2* exon2 including R47H mutation were used for recombination to avoid unwanted splicing described before (Xiang et al., 2018). We also added 66 bases corresponding to the 3xFLAG peptide sequence after the signal peptide to enhance the detection of Trem2 proteins. Insertion of 3xFLAG and human exon2 into the mouse genome was verified by sequencing. However, Trem2 protein with 3xFLAG and human exon2 was not detected with anti-FLAG antibody, although the mRNA was expressed at similar levels as in wt mice (data not shown). The inserted 3xFLAG peptide might affect the processing of the signal peptide or glycosylation at N20 just after the 3xFLAG peptide, leading to unexpected degradation of the protein. We then deleted the 3xFLAG sequence using CRISPR/Cas9 (Fig. 1A) and succeeded in expressing the Trem2 protein whose exon2 was replaced by human exon2 including R47H (designated as *Trem2^hE2R47H^*) (Fig. 1B lane 4). As a control, *Trem2^hE2WT^*mice with human wild-type (WT) exon2 were also generated (Fig. 1A), and equivalent protein expression was confirmed (Fig. 1B lanes 3 and 4). We noticed that the ratio of high molecular weight (40–48 kDa) to low molecular weight (30–33 kDa) of Trem2 was lower in the humanized *Trem2* KI mice (lanes 3 and 4) than in wt mice (lane 1). This might be caused by lower glycosylation efficiency in human exon2 than in mouse exon2, indicating that the humanized *Trem2* (*Trem2^hE2WT^* and *Trem2^hE2R47H^*) are more relevant to human *TREM2* than mouse *Trem2* in glycosylation. Hereafter wt mice were used as a control for *Trem2^KO^* mice, and *Trem2^hE2WT^* as a control for *Trem2^hE2R47H^* and homozygous female mice were analysed.

**Fig. 1.**
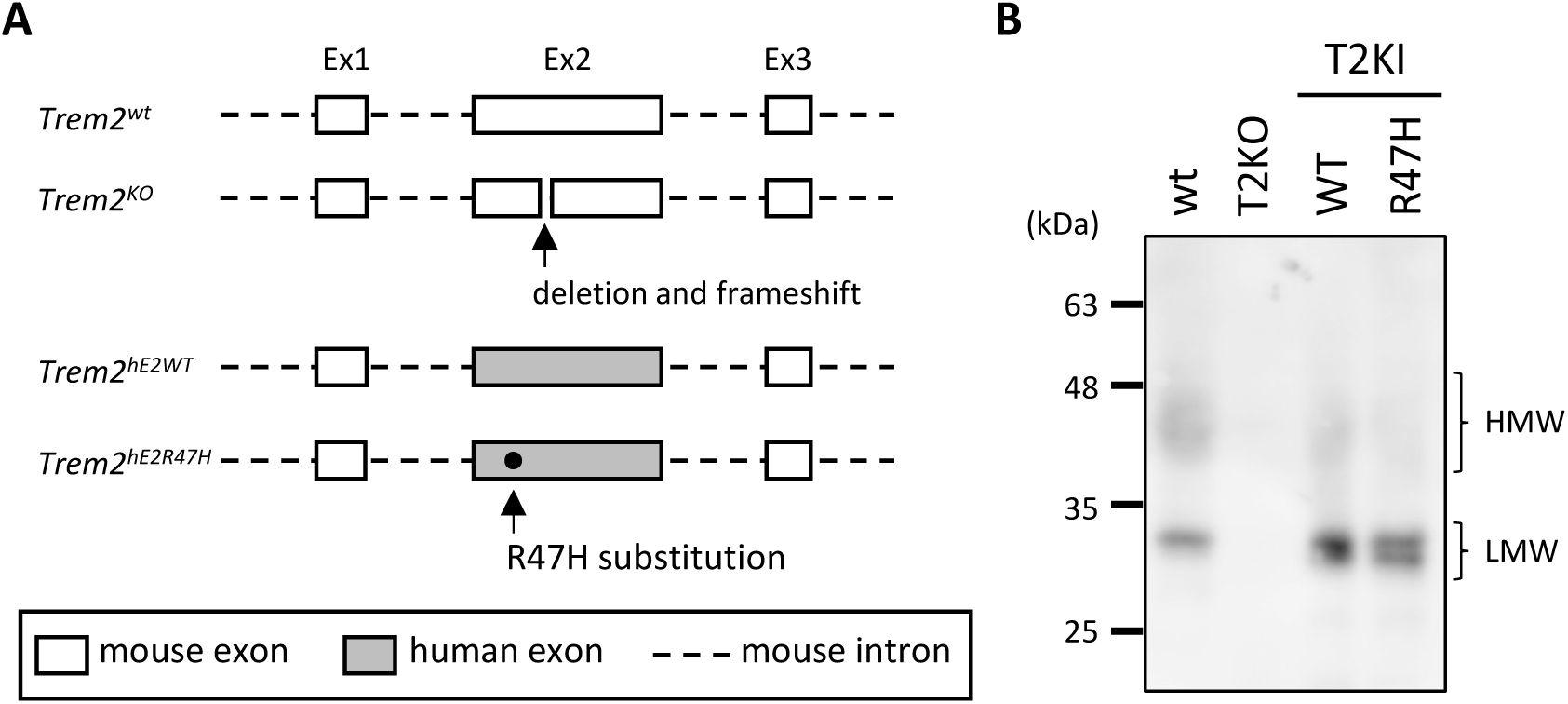
Schematic structure of targeted *Trem2* alleles and expression of Trem2 proteins in *Trem2* KO and *Trem2* KI mice. (A) Schematic representation of *Trem2* alleles (wt, KO, hE2WT, and hE2R47H). Mouse exons and introns are shown as white squares and dashed lines, respectively. Human exon2 is shown by grey squares. (B) Lysates of primary microglia of wt (lane 1), *Trem2^KO^* (lane 2: T2KO), *Trem2^hE2WT^* KI (lane 3: WT) and *Trem2^hE2R47H^*KI mice (lane 4: R47H) were immunoblotted using anti-mouse Trem2 C-terminal antibody. HMW and LMW indicate high and low molecular wights of Trem2 proteins, respectively.

### 3.2. Levels of formic acid-extracted amyloid β 42 were higher in Trem2^KO^ mice but not affected in Trem2^hE2R47H^ mice

To determine whether the genetic modification of *Trem2* alters Aβ pathology in *App^NL-F^* background, we crossed *Trem2^KO^*mice or *Trem2* KI (*Trem2^hE2WT^ or Trem2 ^hE2R47H^*) mice with *App^NL-F^* mice and quantified GuHCl-extracted and subsequently FA-extracted cerebrocortical Aβ42, a dominant amyloid species in the mice. At 18 months, when amyloid plaques are in the growth stage (Saito et al., 2014), GuHCl-extracted Aβ42 levels in *Trem2^KO^* mice were not different from those in wt mice but FA-extracted Aβ42 levels in *Trem2^KO^*mice were almost double (Fig. 2A), suggesting that Trem2 may function to specifically suppress the accumulation of highly fibrillar Aβ42, which can form a core of plaques. *Trem2^KO^* mice at 24 months also had increased amounts of formic acid-extracted Aβ42 compared to wt mice (data not shown). Immunostaining for Aβ and Iba1 (Fig. 2B) revealed that microglia in *Trem2^wt^*x *App^NL-F^* mice clustered around relatively large amyloid plaques (approximately 50 μm in diameter or more) possibly containing highly fibrillar Aβ42 while microglia in *Trem2^KO^* x *App^NL-F^*mice were sparsely distributed, corroborating previous data (Jay et al., 2017, 2015; Meilandt et al., 2020; Wang et al., 2016, 2015; Yuan et al., 2016). Defects in microglial clustering around relatively large amyloid plaques might cause the accumulation of FA-extracted Aβ42 in *Trem2^KO^* mice.

**Fig. 2.**
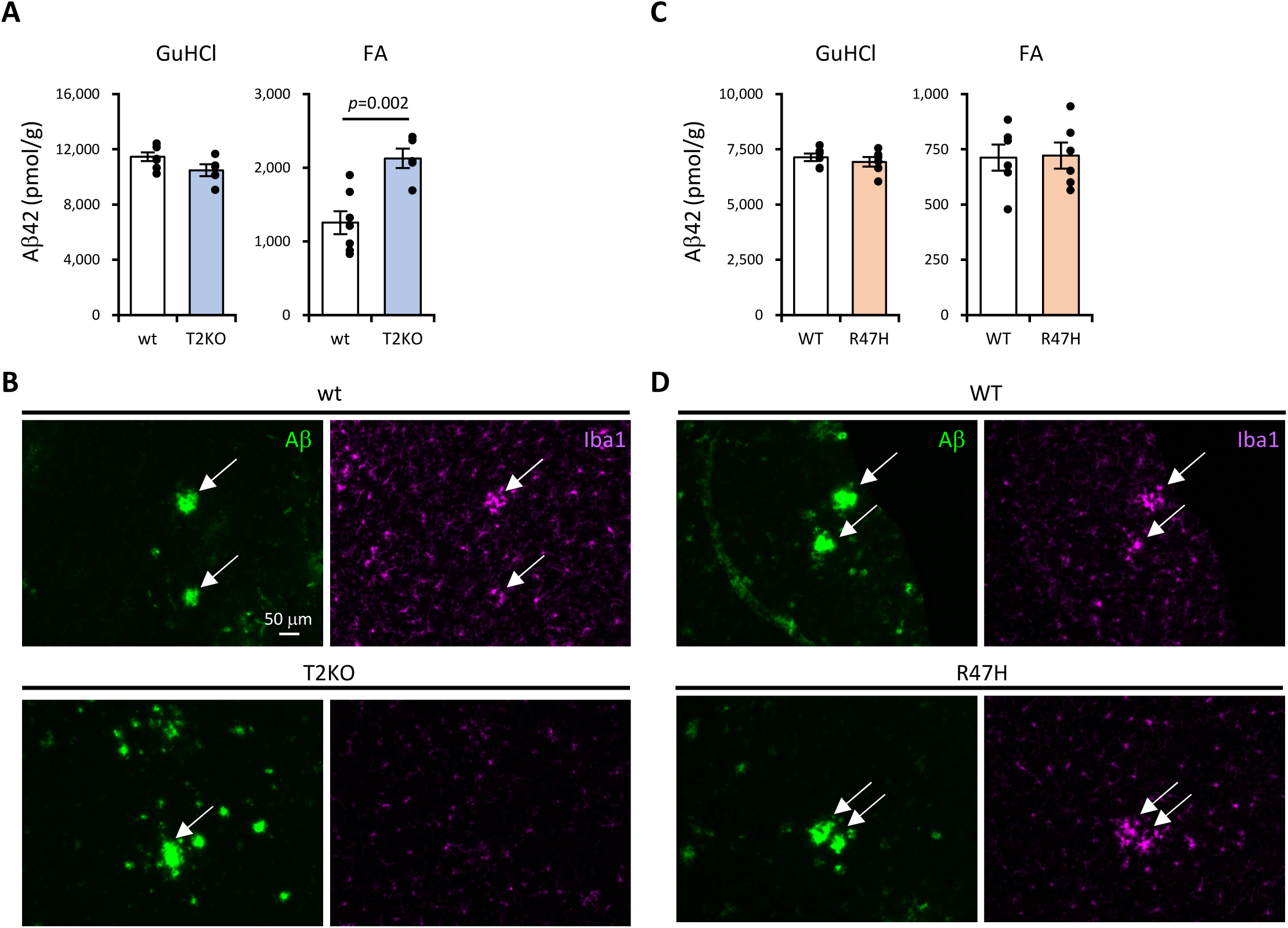
Aβ42 levels in cerebral cortices of 18-month-old *Trem2* KO and *Trem2* KI mice in *App^NL-F^* background (A) and (C) Levels of GuHCl- and FA-extracted Aβ42 were measured using sandwich ELISA. Data are shown as means ± SE (wt, n=7; *Trem2^KO^*, n=5; *Trem2^hE2WT^*, n=6; and *Trem2^hE2R47H^*, n=6). Student t-test was used for statistics. (B) and (D) Representative images of amyloid plaques (green) and Iba1-immunostained microglia (magenta) from brain sections of 18-month-old mice. (A, B) wt and *Trem2^KO^* mice and (C, D) *Trem2^hE2WT^*(WT) and *Trem2^hE2R47H^* (R47H) mice. Images were acquired using 20× objective. Arrows indicate relatively large amyloid plaques and clusters of microglia at the corresponding positions.

However, we did not find any statistically significant differences in Aβ42 levels between *Trem2^hE2WT^* and *Trem2^hE2R47H^*mice at 18 months, either in GuHCl- or in FA-extracted fractions (Fig. 2C). Clustering around relatively large amyloid plaques was apparently not different between these KI mice (Fig. 2D). These results suggest that *Trem2* R47H did not lose, if any, function to keep highly fibrillar Aβ42 at lower levels in contrast to Trem2 deficiency.

### 3.3 Dystrophic neurite formation was enhanced by Trem2 KO, not by Trem2 R47H KI

Damage to axons and dendrites in the vicinity of amyloid plaques, termed dystrophic neurites, is observed in *APP* tg mice and AD patients (Masliah et al., 1996) and genetic deletion of *Trem2* enhances dystrophic neurite formation in *APP* tg mice (Meilandt et al., 2020; Wang et al., 2016; Yuan et al., 2016). We investigated whether dystrophic neurites surrounding amyloid plaques were increased in *Trem2^KO^* x *App^NL-F^*or *Trem2 ^hE2R47H^* x *App^NL-F^* mice by immunostaining cerebrocortical sections for Aβ and Lamp-1, a lysosomal protein enriched in dystrophic neurites. The area of dystrophic neurites surrounding the amyloid plaque was larger in *Trem2^KO^* x *App^NL-F^*mice than in *Trem2^wt^* x *App^NL-F^* mice at 24 months (Fig. 3A). For quantification we normalized the dystrophic neurite area by each amyloid plaque area and compared the ratio between wt and *Trem2^KO^* mice in three categories of amyloid plaque areas (Fig. 3B left). Two-way ANOVA statistical interaction test was performed to determine whether there was a genotype- or plaque area-dependent effect on dystrophic neurite formation. Although no interaction between genotype and plaque area was detected, genotype had a significant effect on dystrophic neurite formation, indicating that *Trem2^KO^*x *App^NL-F^* mice exhibited more dystrophic neurites than *Trem2^wt^*x *App^NL-F^* mice (Fig. 3B left). Actually, *post hoc* test identified a significant increase in the Lamp-1/Aβ area ratio in smaller plaque areas (70–400 μm and 400–800 μm) in *Trem2^KO^* x *App^NL-F^*mice (Fig. 3B). A significant effect of amyloid plaque area on dystrophic neurites was also demonstrated, confirming that smaller, actively growing plaques have a larger halo of dystrophic neurites (Condello et al., 2011; Sadleir et al., 2025, 2022) irrespective of *Trem2* expression. However, *Trem2 ^hE2R47H^* x *App^NL-F^* mice did not display altered dystrophic neurite formation compared to control mice at 24 months (Fig. 3B right) or at 18 months (Fig. 3C). These results clearly indicated that *Trem2^KO^* and *Trem2^hE2R47H^*do not present the same pathological phenotypes.

**Fig. 3.**
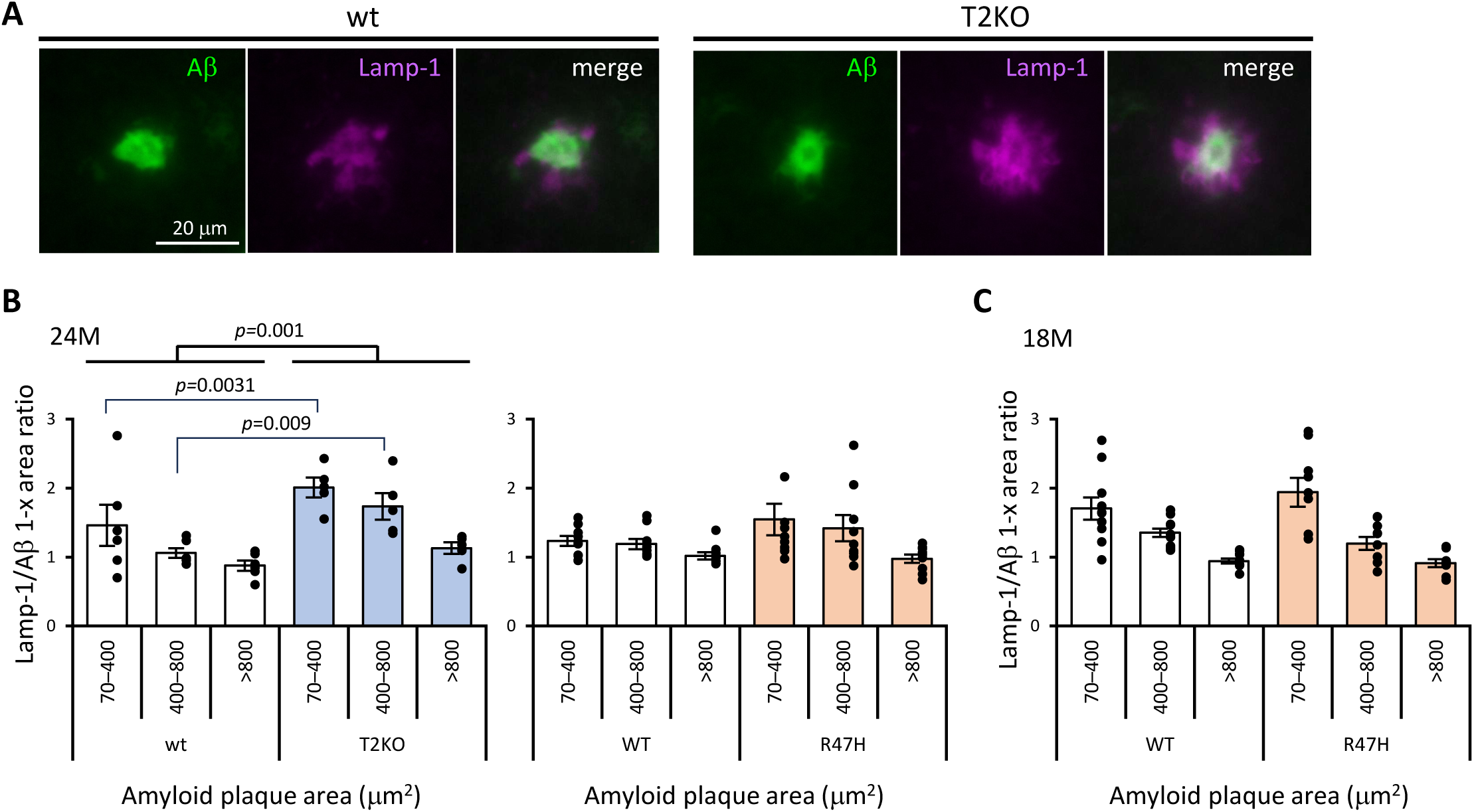
Plaque-associated dystrophic neurites of *Trem2* KO and *Trem2* KI mice in *App^NL-F^*background. (A) Representative images of amyloid plaques (green) and Lamp1-immunostained dystrophic neurites (magenta) from brain sections of 24-month-old wt and *Trem2^KO^* mice. Images were acquired using 100× objective. (B) Ratio of dystrophic neurite and amyloid plaque areas of 24-month-old mice (wt, n=6; *Trem2^KO^*, n=5; *Trem2^hE2WT^*, n=9; and *Trem2^hE2R47H^*, n=9) are represented as means ± SE in each amyloid plaque area (70–400, 400–800, >800 μm^2^). Two-way ANOVA showed a significant main effect between wt and *Trem2^KO^* (F(1, 27) = 12.646; *p* < 0.001). (C) Ratio of dystrophic neurite and amyloid areas of 18 months old mice (*Trem2^hE2WT^*, n=11 and *Trem2^hE2R47H^*, n=9).

### 3.4. Microglial gene expression in Trem2^KO^ and Trem2^hE2R47H^ mice

Since Trem2 is a microglial receptor that regulates gene expression via some signalling pathways in animal models of neurodegenerative disorders (Keren-Shaul et al., 2017; Krasemann et al., 2017), we explored whether the expression levels of microglial genes, including microglial marker and disease-associated microglia (DAM) genes, were altered in *Trem2^KO^*x *App^NL-F^* or *Trem2 ^hE2R47H^* x *App^NL-F^*mice at 18 months. As expected, the levels of *Aif1*, *Tyrobp* and *Cd68* mRNAs were lower in *Trem2^KO^* mice than in wt mice (Fig. 4A left), indicating a failure to transition into an activated microglial state. Next, we first confirmed that *Trem2* mRNAs were expressed at similar levels in both *Trem2* KI mice (Fig. 4B) and then found that mRNA levels of *Aif1*, *Tyrobp* and *Cd68* were comparable between the two groups (Fig. 4A right). These results suggest that *Trem2* R47H still retains the gene regulatory function for these microglial genes in *App^NL-F^* mice.

**Fig. 4.**
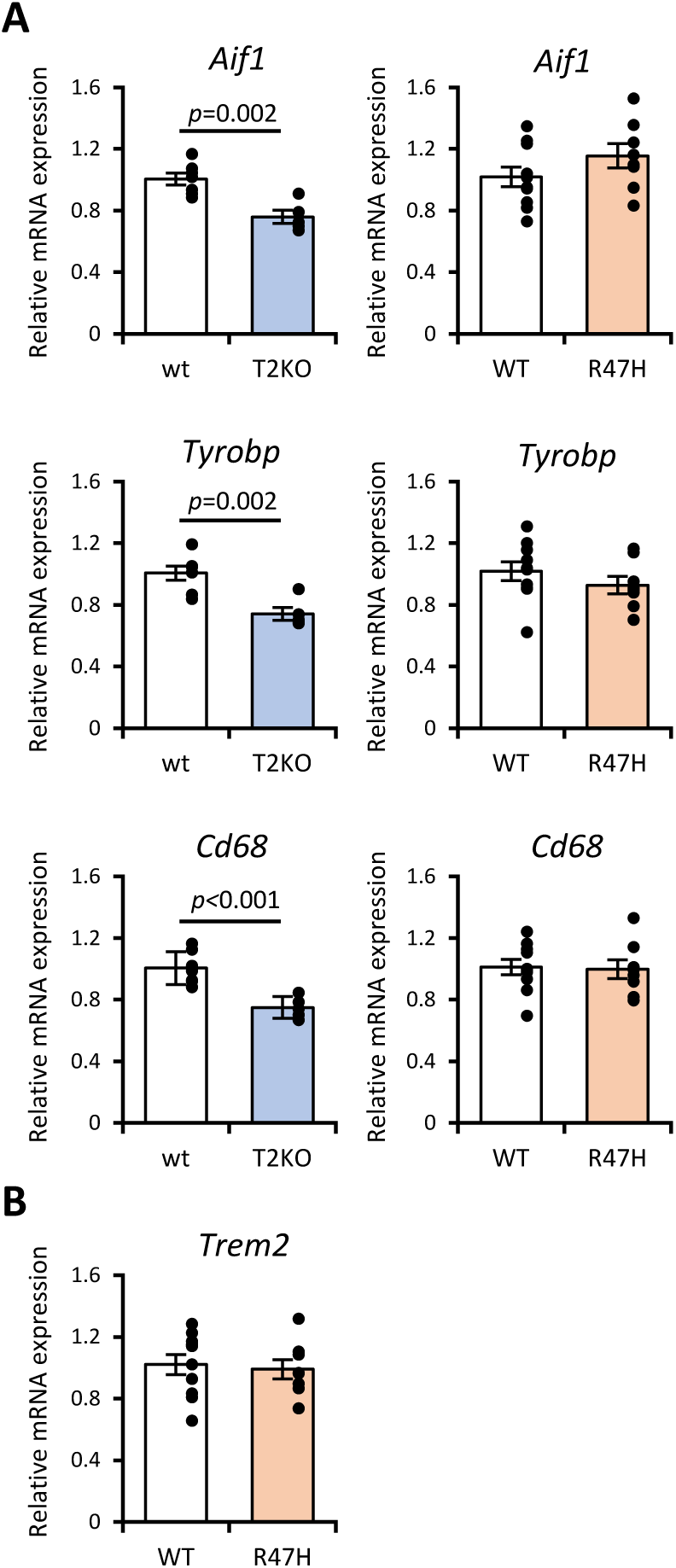
mRNA expression levels of microglial genes in *Trem2* KO and *Trem2* KI mice mRNA levels of microglial genes (A) *Aif1, Tyrobp,* and *Cd68*, and (B) *Trem2* in the cerebral cortices of mice at 18 months were determined by qPCR. Each column represents the mean ± SEM (wt, n=7; *Trem2^KO^*, n=5; *Trem2^hE2WT^*, n=10; *Trem2^hE2R47H^*, n=8). Student t-test was performed and *p*-values are shown where statistical significance was proven.

### 3.5. Transcriptome analysis in Trem2^hE2R47H^ mouse brain

To address gene expression changes by *Trem2* R47H, cerebrocortical total RNA from *Trem2^hE2WT^* x *App^NL-F^*and *Trem2^hE2R47H^* x *App^NL-F^* mice at 18 months were subjected to RNA-seq analysis. We identified 1,092 differentially expressed genes (DEGs) based on the cutoffs of adjusted *p*–value < 0.05. Gene Ontology (GO) (Fig. 5A) and Kyoto Encyclopedia of Genes and Genomes (KEGG) pathway (Fig. 5B) revealed that the DEGs were enriched in ER stress/unfolded protein response and signalling pathways (TNF, Oxytocin, FoxO and MAPK) suggesting that *Trem2* R47H disturbs ER homeostasis and transduces aberrant signals. We selected 13 DEGs with log_2_FC more than 0.25 or less than –0.25 and whose relationships with AD have been reported in at least five papers on the PubMed website (https://pubmed.ncbi.nlm.nih.gov/) (Fig. 5C). These included nine upregulated genes (*Twist1, Tecta, Hdc, Hspbp1, Casr, Ace2, Tgfb1, FosB,* and *Tmem119)* and four downregulated genes (*Gpnmb, Omp, Ada,* and *Pmel*). Moreover, none of the microglial markers, homeostatic or DAM genes (19 genes) were affected by *Trem2* R47H (Fig. 5D), suggesting that *Trem2* R47H may not impair the transition from homeostatic microglia to DAM in *App^NL-F^* mice.

**Fig. 5.**
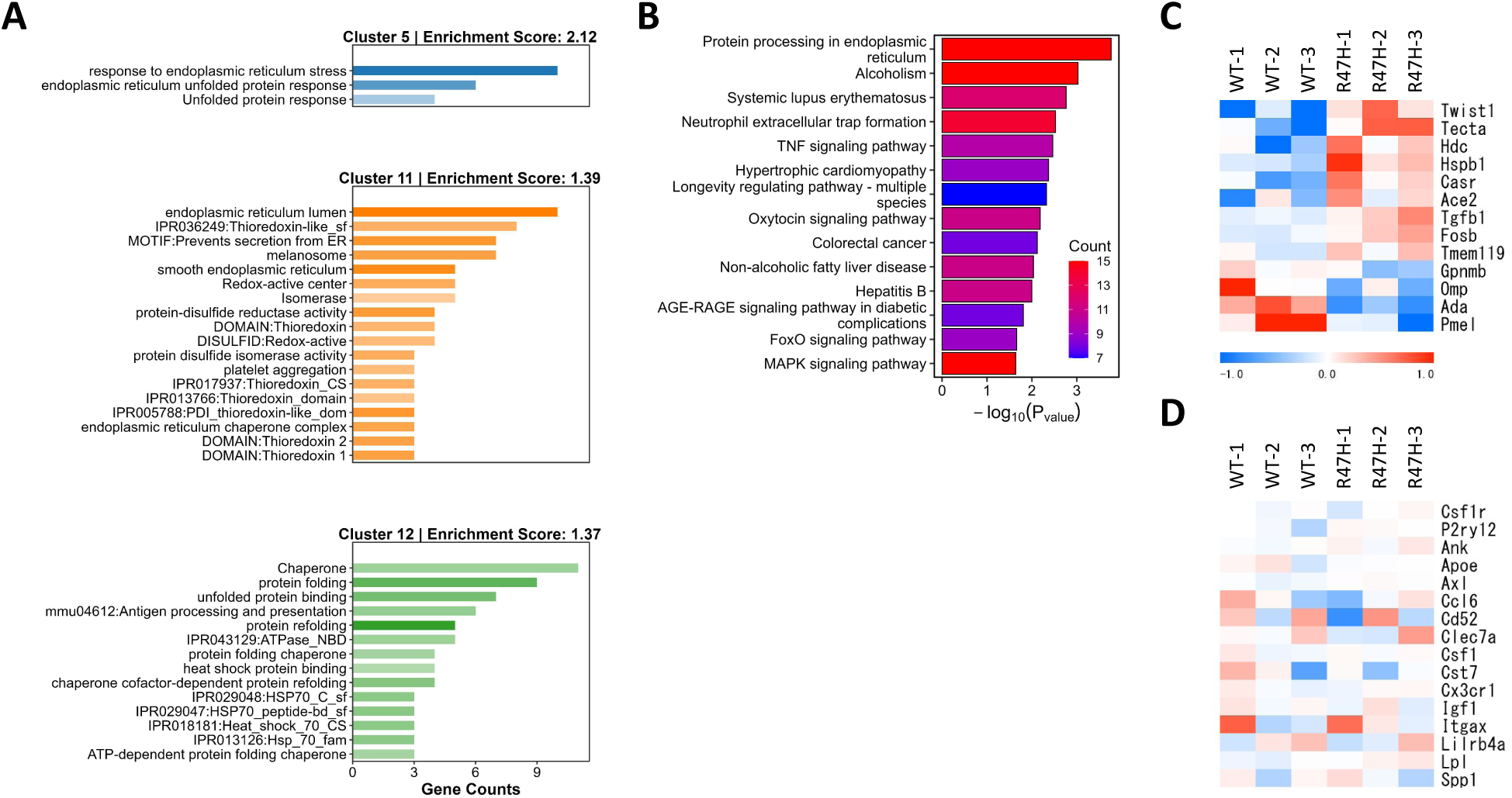
Transcriptional profiling of *Trem2^hE2WT^* and *Trem2^hE2R47H^* mice in *App^NL-F^* background Gene ontology (A) and KEGG pathway (B) of the DEGs between *Trem2^hE2WT^* x *App^NL-F^* and *Trem2^hE2R47H^* x *App^NL-F^* mice. Heatmap of selected DEGs (C) (see text) and microglial markers, homeostatic and DAM genes (D). The color indicates the distance from the median of each row.

### 3.6. Gene expression affected in Trem2 ^hE2R47H^ x App^NL-F^ mice

Among 13 DEGs in Fig. 5C, we selected 4 genes (*Casr, Tmem119, Gpnmb* and *Pmel*) as following reasons. Casr, a member of the G protein-coupled receptor expressed in neurons, is upregulated in AD models (Gardenal et al., 2017) and may induce synaptic injury by binding to Aβ oligomers (Feng et al., 2020). Interestingly, a genetic study has indicated that polymorphisms of *CASR* are associated with AD (Conley et al., 2009). Tmem119 is a marker of homeostatic microglia, which is downregulated in *APP* tg mice (Keren-Shaul et al., 2017; Krasemann et al., 2017) and is involved in the phagocytosis of Aβ (J. Liu et al., 2025). Gpnmb, glycoprotein nonmetastatic melanoma protein B, is one of the DAM genes (Krasemann et al., 2017) and involved in phagocytosis of pathological particles including Aβ (M. Liu et al., 2025). Pmel is homologous to Gpnmb and a substrate of BACE2 (Rochin et al., 2013), although its precise functions in AD remain to be determined. We analyzed mRNA levels of these 4 genes in *Trem2^hE2R47H^* x *App^NL-F^*mice as well as in *Trem2^KO^* x *App^NL-F^* mice. Levels of *Casr* and *Tmem119* were unchanged in *Trem2^hE2R47H^*x *App^NL-F^* mice (Fig. 6A and 6B right) or in *Trem2^KO^*x *App^NL-F^* mice (Fig. 6A and 6B left). *Gpnmb* levels tended to be lower in both *Trem2^hE2R47H^* x *App^NL-F^*mice (Fig. 6C right) and in *Trem2^KO^* x *App^NL-F^* mice (Fig. 6C left) but the difference from the control did not reach statistical significance. Finally, *Pmel* levels were significantly lower in *Trem2^hE2R47H^*x *App^NL-F^* mice (Fig. 6D right) and tended to be lower in *Trem2^KO^* x *App^NL-F^* mice (Fig. 6D left).

**Fig. 6.**
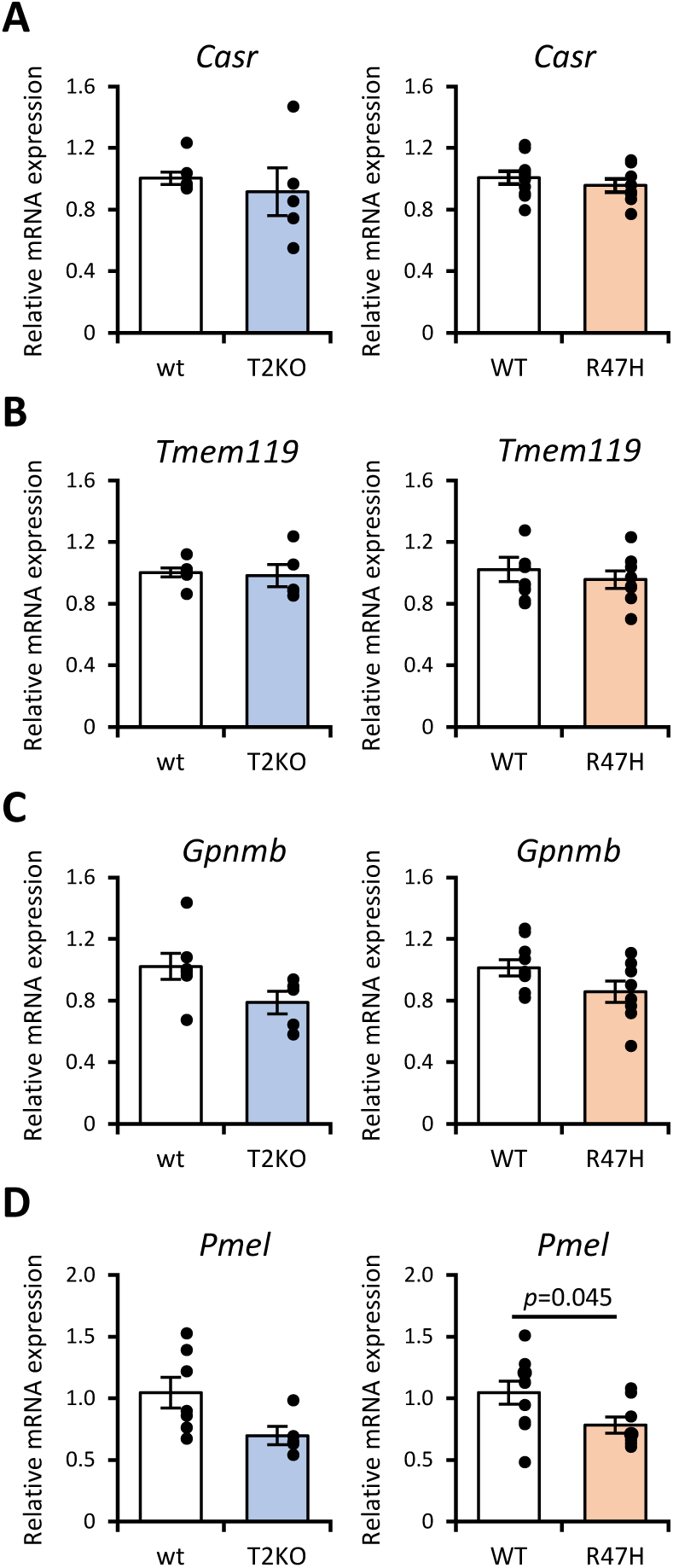
mRNA expression levels of the DEGs in *Trem2* KO and *Trem2* KI mice mRNA levels of the DEGs identified by RNA-seq analysis (*Casr, Tmem119, Pmel,* and *Gpnmb*) in the cerebral cortices of mice at 18 months were determined by qPCR. Each column represents the mean ± SEM (wt, n=7; *Trem2^KO^*, n=5; *Trem2^hE2WT^*, n=10; *Trem2^hE2R47H^*, n=8). Student t-test was performed and *p*-values are shown where statistical significance was proven.

## 4. Discussion

The present study comprehensively evaluated the functional impact of the TREM2 R47H variant in the context of Aβ pathology. The use of the aged *App^NL-F^*mouse model represents a major strength of this study. Because this model avoids APP overexpression, and Aβ42 accumulation continues gradually until at least 24 months, it provides a physiologically relevant context for evaluating microglial function and Aβ pathology in late-onset AD. By directly comparing *Trem2* R47H KI mice with *Trem2* KO mice, we demonstrated that the R47H variant confers a minimal loss-of-function phenotype, resulting in only subtle alterations in gene expression. These findings clarify the degree to which this human AD risk variant disrupts microglial responses and highlight the importance of *Trem2* R47H KI mice in analyzing mechanism by which TREM2 enhances the incidence of AD.

To analyze the pathological mechanism by which incidence of AD is increased by *TREM2* R47H, four KI mice had been produced (Cheng et al., 2018; Cheng-Hathaway et al., 2018; Xiang et al., 2016). However, three lines out of four showed unexpected cryptic mRNA splicing due to the introduced mutations corresponding to R47H and synonymous silent codon changes and subsequent reduction of full-length Trem2 mRNA and protein expression. Protein levels of Trem2 R47H in the KI mouse line made by Cheng et al. were preserved but AD pathology in the mice have not been analyzed (Cheng et al., 2018). Since Xiang et al. demonstrated exon2 of human TREM2 containing R47H does not induce cryptic splicing (Xiang et al., 2018), we replaced mouse exon2 with human exon2 containing the R47H mutation and successfully produced *Trem2* R47H KI mice (Fig. 1A) without a reduction in Trem2 mRNA levels (Fig. 4B) and protein expression (Fig. 1B). Recently Sayed et al. knocked-in human *TREM2* R47H cDNA into the mouse Trem2 locus (Sayed et al., 2021). Although they confirmed that TREM2 R47H protein expression in the mice is comparable with that in WT TREM2 cDNA KI mice, aging- and pathology-dependent spatial expression of TREM2 in the brain might not be reproduced because the KI mice have no introns that may regulate gene expression. Tran et al. produced *Trem2* R47H KI mice without cryptic splicing and analyzed AD pathology in the mice (see details below) (Tran et al., 2023). On the other hand, our *Trem2* KI mice contain human exon2, the longest exon of *Trem2* which encodes more than half of Trem2 protein (117 amino acids out of total 227 amino acids). In addition, since exon2 contains an Ig-like v-type domain and two N-glycosylation sites, which play a role in ligand binding (Kober et al., 2016; Sudom et al., 2018) and trafficking (Shirotani et al., 2022) respectively, exon2-humanized *Trem2* KI mice might be more suitable for understanding human pathology. Functional difference among species have been observed in the case of the other AD risk gene ApoE, as knock-in of human ApoE isoforms resulted in a delay of Aβ deposition relative to murine ApoE (Fryer et al., 2005).

Amounts of FA-extracted Aβ42 were increased in *Trem2^KO^*mice at 18 months (Fig. 2A) and 24 months (data not shown). In addition, deletion of *Trem2* abrogated microglial clustering around relatively large amyloid plaques (Fig. 2B). These results suggest that Trem2 plays a key role in microglial clustering around relatively large amyloid plaques and phagocytosing fibrillar Aβ42 (Kleinberger et al., 2014). It is also possible that Trem2 inhibits Aβ42 fibrilization by binding to Aβ42 as suggested previously (Belsare et al., 2022; Kober et al., 2021; Vilalta et al., 2021).

However, amounts of FA-extracted Aβ42 in *Trem2^hE2R47H^*mice were not different from those in control *Trem2^hE2WT^* mice (Fig. 2C). This suggests that the arginine residue at the 47th position might not be involved in suppressing the formation of highly fibrillar Aβ42. Das et al. also reported that amyloid plaque levels are not changed in the human *TREM2* R47H cDNA KI mice crossed with *App^NL-G-F^*(Das et al., 2023). Tran et al. also reported that in *Trem2^R47H^* x 5xFAD mice amounts of FA-extracted Aβ42 in cerebral cortex were not altered at 4 months (plaque growth stage in 5xFAD mice) but increased at 12 months (plaque plateaus stage) (Tran et al., 2023). Further analysis of Aβ42 at various ages might provide insight into the functions of *Trem2* R47H during aging. Taken together, our findings demonstrate that the R47H variant does not completely recapitulate TREM2 deficiency.

Dystrophic neurites around amyloid plaques were exacerbated by *Trem2* KO in *App^NL-F^* mice (Fig. 3A and 3B), consistent with previous studies in *APP* tg mice (Meilandt et al., 2020; Wang et al., 2016; Yuan et al., 2016). The increased dystrophic neurites might be caused by increased levels of highly fibrillar Aβ42 in *Trem2^KO^* x *App^NL-F^*mice (Fig. 2A), supporting the notion that the transition from a diffuse to β-sheet-rich filamentous conformation of amyloid plaques is a critical event that leads to dystrophy (Yuan et al., 2016). In fact, cored or coarse-grained plaques induce more dystrophic neurites in their vicinity than diffuse plaques (Koutarapu et al., 2025). It is also possible that microglia in *Trem2^KO^* mice cannot suppress the contact between amyloid plaques and axons/dendrites, leading to enhanced neuritic dystrophy. We noticed that as plaques grew larger, the difference in axonal dystrophy between wt and *Trem2^KO^* diminished (Fig. 3B left, amyloid plaque area >800 mm^2^). The degree of axonal dystrophy around larger plaques might have reached a plateau, preventing microglia from suppressing dystrophic neurite formation.

In contrast, we could not detect differences in dystrophic neurite formation between *Trem2^hE2WT^* and *Trem2^hE2R47H^*mice at either 18 or 24 months (Fig. 3B right and Fig. 3C). This might be because the FA-extracted Aβ42 levels were unchanged by the *Trem2* R47H mutation (Fig. 2C). These results also corroborate that the R47H variant has a weak and hypomorphic phenotype. Tran et al. reported that dystrophic neurites are enhanced in *Trem2^R47H^*x 5xFAD mice at 4 months (Tran et al., 2023). The difference from our result might be due to the different mouse models used.

mRNA levels of microglial marker and DAM genes were lower in *Trem2^KO^*x *App^NL-F^* mice than in *Trem2^wt^* x *App^NL-F^*mice (Fig. 4A left), suggesting that Trem2 is responsible for the upregulation of microglial genes in *App^NL-F^* mice, as shown in *APP* tg mice (Keren-Shaul et al., 2017; Krasemann et al., 2017). We also found that these gene expression changes were not observed in *Trem2^hE2R47H^* x *App^NL-F^*mice (Fig. 4A right), suggesting that the R47 residue is not critical for signal transduction that induces these microglial genes. Consistently, Tran et al. reported that most DAM genes were not affected in their *Trem2^R47H^* x 5xFAD mice (Tran et al., 2023).

RNA-seq and differential gene expression analyses suggested that genes involved in ER stress/unfolded protein response (Fig. 5A) and signalling pathways of TNF, Oxytocin, FoxO and MAPK (Fig. 5B) were altered in *Trem2^hE2R47H^* x *App^NL-F^*mice. Since TREM2 R47H protein accumulates in aberrant glycoforms due to prolonged retention within the ER (Park et al., 2017, 2015), the R47H variant could increase ER stress/unfolded protein response under stress-sensitized conditions. Interestingly, ER stress and unfolded protein response perturb the fidelity and pattern of N-linked glycosylation (Phoomak et al., 2021; Wong et al., 2018; Yung et al., 2023). In addition, MAPK signalling alters glycan structures by regulating the expression of glycosyltransferases (Pan et al., 2024; Park et al., 2012; Wu et al., 2024). As we have demonstrated that glycosylation of TREM2 is required for trafficking of the protein to the cell surface as well as for signal transduction (Shirotani et al., 2022), altered ER stress/unfolded protein response and MAPK signalling might cause aberrant cellular localization of Trem2 protein and affect its signalling activity, which may contribute to AD development.

Among the 13 DEGs (Fig. 5C), we confirmed that *Gpnmb* and *Pmel* showed reduced expression or tendency in *Trem2^hE2R47H^* x *App ^NL-F^*mice (Fig. 6C and 6D). Gpnmb, glycoprotein nonmetastatic melanoma protein B, is expressed on plasma membrane, endosome and lysosome of microglia and a risk factor of Parkinson’s disease (Diaz-Ortiz et al., 2022). Interestingly, Gpnmb internalizes fibrillar α-synuclein into synapses (Diaz-Ortiz et al., 2022). Recently Liu et al. showed that Gpnmb is induced by pathological particles including Aβ and participates in phagocytosis/degradation of them by association with lysosomal ATP6V1A (M. Liu et al., 2025). Therefore, decreased expression of Gpnmb by *Trem2* R47H may lead to the accumulation of Aβ in AD patients. Pmel is expressed on melanocytes of skin and forms physiological amyloids that play a central role in melanosome morphogenesis and pigmentation. Our qPCR detected *Pmel* in the mouse cerebral cortex. Moreover, Human protein atlas database (https://www.proteinatlas.org/) suggests the expression of *Pmel* in microglia. Further investigation of microglial Pmel may identify novel therapeutic molecular targets.

Although we identified 13 DEGs that may be related to AD pathogenesis (Fig. 5C), these genes were not affected in *Trem2^R47H^* x 5xFAD mice (Tran et al., 2023). Moreover, DEGs in *Trem2^R47H^* x 5xFAD mice, except for one, were not altered in our *Trem2^hE2R47H^* x *App ^NL-F^* mice. Only the expression change of *Mid1* was directionally consistent in both *Trem2^hE2R47H^* x *App^NL-F^*and *Trem2^R47H^* x 5xFAD mice. Mid1, expressed in neurons and astrocytes, was upregulated 1.57-fold at 18 months in our *Trem2^hE2R47H^*x *App ^NL-F^* mice, while 1.31-fold at 4 months and 1.77-fold at 12 months in *Trem2*^R47H^ x 5xFAD mice (Tran et al., 2023). Because Mid1 is a ubiquitin E3 ligase for immunoglobulin-binding protein 1 and regulates the levels of protein phosphatase PP2A, which dephosphorylates tau (Watkins et al., 2012), upregulation of Mid1 may cause degradation of PP2A and a subsequent increase in tau phosphorylation, contributing to AD pathogenesis.

This study has several limitations that should be acknowledged. First, this study focused on bulk RNA-seq, which may obscure cell-type–specific transcriptional changes. Isolation of microglia and single-cell or spatial transcriptomics could provide deeper insights into how R47H alters microglial response. Second, we did not assess functional outcomes such as phagocytic capacity or plaque compaction at the ultrastructural level. Finally, although the *App^NL-F^* model recapitulates key aspects of Aβ pathology, it does not capture tau pathology or neurodegeneration, which may interact with TREM2 signaling in additional ways.

In conclusion, our findings demonstrate that the TREM2 R47H variant produces only mild impairments in microglial responses to Aβ pathology, which is distinct from the severe deficits caused by complete TREM2 loss. This is reasonable considering that TREM2 is a risk gene of AD, not a causal gene. Detection of clear effects of TREM2 R47H on AD pathology would be difficult in *APP* tg, *App^NL-G-F^*, and even *App^NL-F^* KI mice, where Aβ accumulation velocity is still higher than that in humans. Although pathological phenotypes are not easily detected in *Trem2^hE2R47H^*x *App ^NL-F^* mice, changes in gene expression may occur at the earlier phases. We are currently planning transcriptome analysis at 6 or 12 months. Moreover, if we use animal models that show slower Aβ accumulation, such as *App^NL^* KI mice (Saito et al., 2014), the phenotypes of *Trem2^hE2R47H^* mice might be detected. Taken together, *Trem2^hE2R47H^*x *App^NL-F^* mouse model established here indicates that the R47H variant has subtle but reasonable effects on AD pathology and microglial gene expression but it should contribute to the identification of novel AD mechanisms and the development of drugs.

## Abbreviations

Aβ: amyloid β peptide
AD: Alzheimer’s disease
ANOVA: analysis of variance
APP: amyloid-β precursor protein
ER: endoplasmic reticulum
FA: formic acid
GO: Gene ontology
GuHCl: guanidine hydrochloride
KEGG: Kyoto Encyclopedia of Genes and Genomes
KI: knock-in
KO: knockout
RNA-seq: RNA sequencing
tg: transgenic
TREM2: triggering receptor expressed on myeloid cells 2
WT: human wild-type
wt: mouse wild-type

## CRediT authorship contribution statement

Keiro Shirotani and Nobuhisa Iwata: Conceptualization, Data curation, Investigation, Methodology, Writing – original draft, Writing – review & editing, Validation, Visualization, Funding acquisition. Daisuke Hatta: Writing – review & editing, Data curation. Kaori Watanabe: Investigation, Methodology. Takashi Saito and Takaomi C. Saido: Writing – review & editing, Resources.

## Consent for publication

All authors consented to submit the manuscript to the journal.

## Declaration of generative AI and AI-assisted technologies in the manuscript preparation process

During the preparation of this work, the author(s) used Paperpal (Editage) to improve the language and readability. After using this tool, the author(s) reviewed and edited the content as needed and took full responsibility for the content of the published article.

## Acknowledgements

We thank Aki Noda, Gaku Shizuki, Anna Ishimoto and Takaaki Teranishi for assistance of the experiments.

## Data availability

The data are available from the corresponding author upon reasonable request.

